# Association of Polygenic Risk Scores for Multiple Cancers in a Phenome-wide Study: Results from The Michigan Genomics Initiative

**DOI:** 10.1101/205021

**Authors:** Lars G. Fritsche, Stephen B. Gruber, Zhenke Wu, Ellen M. Schmidt, Matthew Zawistowski, Stephanie E. Moser, Victoria M. Blanc, Chad M. Brummett, Sachin Kheterpal, Gonçalo R. Abecasis, Bhramar Mukherjee

**Affiliations:** Department of Biostatistics, University of Michigan School of Public Health, Ann Arbor, MI 48109, USA.; Center for Statistical Genetics, University of Michigan School of Public Health, Ann Arbor, MI 48109, USA.; K.G. Jebsen Center for Genetic Epidemiology, Department of Public Health and Nursing, Norwegian University of Science and Technology, 7491 Trondheim, Sør-Trøndelag, Norway.; USC Norris Comprehensive Cancer Center, Los Angeles, CA 90033, USA.; Michigan Institute for Data Science, University of Michigan, Ann Arbor, MI 48109, USA.; Division of Pain Medicine, Department of Anesthesiology, University of Michigan Medical School, Ann Arbor, MI 48109, USA.; Central Biorepository, University of Michigan Medical School, Ann Arbor, MI 48109, USA.; Institute for Healthcare Policy and Innovation, University of Michigan, Ann Arbor, MI 48109, USA.; Department of Epidemiology, University of Michigan School of Public Health, Ann Arbor, MI 48109, USA.; University of Michigan Comprehensive Cancer Center, University of Michigan, Ann Arbor, MI 48109, USA.; Present Address: Department of Biostatistics and Epidemiology, University of Michigan School of Public Health, 1415 Washington Heights, SPH 1 Room 4619, Ann Arbor, MI 48109, USA.

## Abstract

Health systems are stewards of patient electronic health record (EHR) data with extraordinarily rich depth and breadth, reflecting thousands of diagnoses and exposures. Measures of genomic variation integrated with EHRs offer a potential strategy to accurately stratify patients for risk profiling and discover new relationships between diagnoses and genomes. The objective of this study was to evaluate whether Polygenic Risk Scores (PRS) for common cancers are associated with multiple phenotypes in a Phenome-wide Association Study (PheWAS) conducted in 28,260 unrelated, genotyped patients of recent European ancestry who consented to participate in the Michigan Genomics Initiative, a longitudinal biorepository effort within Michigan Medicine. PRS for 12 cancer traits were calculated using summary statistics from the NHGRI-EBI catalog. A total of 1,711 synthetic case-control studies was used for PheWAS analyses. There were 13,490 (47.7%) patients with at least one cancer diagnosis in this study sample. PRSs exhibited strong association for several cancer traits they were designed for including female breast cancer, prostate cancer, melanoma, basal cell carcinoma, squamous cell carcinoma and thyroid cancer. Phenome-wide significant associations were observed between PRS and many non-cancer diagnoses. To differentiate PRS associations driven by the primary trait from associations arising through shared genetic risk profiles, the idea of “exclusion PRS PheWAS” was introduced. This approach led to phenome-wide significant associations between a lower risk for hypothyroidism in patients with high thyroid cancer PRS and a higher risk for actinic keratosis in patients with high squamous cell carcinoma PRS after removing all cases of the primary cancer trait. Further analysis of temporal order of the diagnoses improved our understanding of these secondary associations. This is the first comprehensive PheWAS study using PRS instead of a single variant.

## Introduction

In the past decade, genome-wide association studies (GWAS) using single nucleotide polymorphisms (SNPs) led to discovery of many common disease susceptibility loci ^1-3^ An alternative agnostic way of exploring gene-disease association is through phenome-wide association studies (PheWAS) ^4-6^ PheWAS enable simultaneous exploration of the association between genetic variants and a broad spectrum of physiological/clinical phenotypes. To explore the joint genome × phenome landscape, one needs access to both Electronic Health Records (EHRs) and GWAS data. The promise and potential of these studies have recently been illustrated by the electronic Medical Records and Genomics (eMERGE) network ^7; 8^. Beyond genetic associations, EHR has enabled discovery of new associations between disease and secondary effects of drugs or blood biomarker levels ^9-11^.

PheWAS have been used to both replicate known genetic-phenotypic associations and to discover new consequences for disease associated variants. PheWAS use computable phenotypes derived from EHR databases. Traditional PheWAS have used International Classification of Disease (ICD) codes to define a set of computable phenotypes or “PheWAS codes” defined and validated by experts using a combination of ICD codes ^12^ Standard PheWAS have primarily focused on correlating genetic variants, **one at a time**, to a spectrum of phenotypes. When each variant is associated with a small effect size, these studies can only provide limited insight. For this reason, many areas of genetics now use ensembles of variants that cumulatively explain substantial variation in disease risk ^13-15^. For example, PRS constructed from multiple GWAS identified loci that have been proposed for cancer screening, risk prediction and risk stratification ^16-19^.

In this paper, we introduce the new concept of exploring PRS in a PheWAS setting instead of a traditional PheWAS that considers single variants, one at a time. We focus on cancer traits while constructing the PRS. We construct PRS for multiple cancers including some of the most common groupings of cancers in the United States: prostate cancer (PCa, MIM: 176807), breast cancer (MIM: 114480), colorectal cancer (MIM: 114500), lung cancer (MIM: 211980), melanoma of skin (MIM: 155601) and basal cell carcinoma (MIM: 614740) and correlate them with PheWAS codes.

Our study is based on the Michigan Genomics Initiative (MGI) launched in 2012, a biorepository effort to create a longitudinal cohort of participants in Michigan Medicine. MGI enrolled participants undergoing anesthesia prior to a surgery or diagnostic procedure, creating a patient community with genome-wide data, electronic health information, and permission for follow-up and re-contact in future studies. Our current analysis of 28,260 patients in MGI indicates that 47.7% of these patients have at least one current or previous neoplasm diagnosis (excluding benign neoplasms). This presents a unique opportunity to study multiple cancer outcomes leveraging both EHR and genomic data in MGI.

At the same time, this enrichment of cancer patients in MGI highlights some of the special features of the sampling frame for the study and source population. Because of the self-selective, consent-based nature of MGI patient enrollment, the sample selection mechanism is non-probabilistic, that is, the probability of a sampling unit being included in the study is not pre-determined. The MGI sample is enriched for neoplasm diagnoses which could be related to the fact that many surgeries are diagnostic procedures are specifically related to cancer treatment and screening (e.g., colonoscopy, skin biopsies). Cancer patients undergo surgery more frequently than the general population and frequently choose an academic medical center for diagnostic and/or interventional procedures. The analytic framework presented in this paper conducts careful sensitivity analysis for protecting our inference against such selection biases, unbalanced case-control ratios, and phenotypic enrichment.

There are several innovative aspects to our study. Our study represents the first comprehensive PheWAS focused on using **PRS in a cancer-enriched cohort** accrued in an academic health center. Our study is also the first PheWAS focused **on cancer**. Our results demonstrate PRS, a summary score constructed based on results of large population-based GWAS, can be potentially useful for cancer risk stratification among patients in an academic medical center. We also note that when a PRS-based PheWAS leads to the association of a cancer-specific PRS (e.g., prostate cancer PRS) with other secondary related phenotypes (e.g., erectile dysfunction or urinary incontinence), these findings may require careful consideration. We observe that many of these secondary associations are often driven by the primary cancer diagnosis. We introduce the notion of “exclusion PRS PheWAS” to detect independent secondary associations that have shared genetic etiology. We extract the temporal order of diagnoses from the EHR to shed further insight into these secondary associations.

## Subjects and Methods

### MGI cohort

Participants were recruited through Michigan Medicine health system while awaiting diagnostic or interventional procedures either during a preoperative visit prior to the procedure or on the day of procedure that required anesthesia. Opt-in written informed consent is obtained. In addition to coded biosamples and protected secure health information, participants understand that all EHR, claims, and national data sources linkable to the participant may be incorporated into the MGI databank. Each participant donates a blood sample for genetic analysis, undergoes baseline vital signs and a comprehensive history and physical, and completes validated self-report measures of pain, mood and function, including NIH Patient Reported Outcomes Measurement Information System (PROMIS) measures. Data were collected according to Declaration of Helsinki principles. Study participants provided written informed consent, and protocols were reviewed and approved by local ethics committees (IRB ID HUM00099605). In the current study, we report results obtained from 28,260 genotyped samples of European ancestry with available integrated EHR data (see summary characteristics of the cohort in **Table 1**).

**Table 1.** Demographics and clinical characteristics of the final analytic data set

| Characteristic | Analytic Data Set |
| --- | --- |
| N | 28,260 |
| Females, N (%) | 15,113 (53.5%) |
| Mean Age, years (S.D.) | 54.1 (15.9) |
| Total number of ICD9 code days | 3.5 million |
| Number of unique ICD9 codes | 10,322 |
| Median number of visits per participant | 23 |
| Median days between first and last visit | 1,265 |
| Median ICD9 code days per participant | 28 |

### Genotyping and Sample Quality Control (QC)

DNA from 37,412 blood samples was genotyped on two batches of customized Illumina HumanCoreExome v12.1 bead arrays (“UM_HUNT_Biobank_11788091_A1” [N = 21,207] and “UM_HUNT_Biobank_v1-1_20006200_A” [N = 16,205]) that in addition to standard genome-wide tagging SNPs (~N=240,000) and exomic variants (N=~280,000) contained about 70,000 additional custom content variants, e.g. candidate variants from GWAS experiments, nonsense and missense variants from sequencing studies, ancestry informative markers, and Neanderthal variants. Genotype analysis was performed with Illumina GenomeStudio (module 1.9.4, algorithm GenTrain 2.0). After initial clustering, variant cluster boundaries were re-defined in a second run using only individuals with call rate of at least 99% and genotyped the remaining samples afterwards.

We excluded samples with: (1) call rate <99%, (2) estimated contamination >2.5 % (BAF Regress)^20^, (3) large chromosomal copy number variants (single chromosome with missingness >= five times larger than other chromosomes), (4) lower call rate than its technical duplicate or twin, (5) gonosomal constellations other than XX and XY, or (6) whose inferred sex did not match the reported gender. We excluded variants if: (1) their probes could not be perfectly mapped or mapped perfectly to multiple position in the human genome assembly (Genome Reference Consortium Human genome build 37 and revised Cambridge Reference Sequence of the human mitochondrial DNA; BLAT) ^21^, (2) they showed deviations from Hardy Weinberg equilibrium in European ancestry samples (P<0.0001), (3) had a call rate <99%, (4) another variant with higher call rate assayed the same variant or (5) if the allele frequency differences between the two array versions within unrelated, European ancestry samples had a P-value < 0.001 (PLINK v1.90)^22^. After quality control, 392,323 polymorphic variants remained.

Before preparing the final analytical data set, we reduced the data to 33,028 samples for which we had complete age and ICD9 data available. Next, we estimated pairwise kinship with the software KING ^23^ and limited further analysis to a subset that contained no pairs of individuals with a 1st‐ or 2nd-degree relationship. We inferred recent ancestry by projecting all genotyped samples into the space of the principal components of the Human Genome Diversity Project reference panel using PLINK (938 unrelated individuals) ^24; 25^. We limited the principal component analysis to variants that were shared between the HGDP reference and the MGI data, had a minor allele frequency >1%, and remained after LD pruning (r^2^<0.5; PLINK). Samples of recent European ancestry (~90% of participants) were defined as samples that fell into a circle around the center of the European HGDP populations in the PC1 versus PC2 space, whereas the circle’s radius was set to 1/8 of the distance between the center of the European HGDP populations and the centroid of the centers of the European, East Asian and Sub-Saharan populations (**Figure S1**). Principal components were stored and used for further association tests. After quality control, 28,260 unrelated, genotyped individuals of recent European ancestry with age and ICD9 data remained for further analysis.

### Phasing and Genotype Imputation

We imputed genotypes of the Haplotype Reference Consortium using the Michigan Imputation Server ^26^ and filtered poorly imputed variants with R^2^<0.3 and/or minor allele frequency (MAF) < 0.1% resulting in over 17 million imputed variants available after quality control and filtering. The obtained accuracy for imputed variants, i.e. the average empirical R^2^ values for different MAF frequency bins, was: 0.89 (0.1%<=MAF≤0.5%), 0.94 (0.5%<MAF≤5%), and 0.96 (MAF>5%).

### Phenome Generation

We extracted the ICD9 data for 28,260 unrelated, genotyped individuals of recent European ancestry and mapped a total of 3.5 million ICD9 codes to PheWAS codes (PheWAS translation table version 1.2)^12^. The ICD9 codes (10,322 unique ICD9 codes) were aggregated to PheWAS traits using the PheWAS R package ^12^. Cases for a given PheWAS code were defined if an individual had at least one assignment of that PheWAS code in their record. The remaining individuals that did not have overlapping PheWAS codes that are a part of the exclusion criteria were considered as controls. A total of 1,857 case control studies were generated of which 1,711 with ≥20 cases were used for further analyses (see **Table S1**; phenotypes with < 50 cases were coded as “<50”).

### GWAS Catalog SNP Extraction and Construction of PRS

We downloaded previously reported GWAS variants from the NHGRI-EBI Catalog (file date: June 31, 2017) ^27; 28^. None of the discovery studies included in the catalog used any subset of the MGI cohort. This is primarily because MGI started recruiting in 2012 and the genotype data only became available recently. Variant positions were converted to GRCh37 using variant IDs from dbSNP version 144 (UCSC genome browser) after updating outdated dbSNP IDs to their merged dbSNP IDs. Entries with missing risk alleles, risk allele frequencies, or odds ratios were excluded. We corrected alleles of non-ambiguous SNPs to the forward strand of the genomic reference sequence so that the reported risk allele matched one of the alleles found at the corresponding position in the 1000 Genomes Project genotype data. We only included entries with broad European ancestry (as reported by the NHGRI-EBI Catalog). To allow an additional quality control check, we compared the reported risk allele frequencies (RAF) in controls with the frequencies of the 503 European samples of the 1000 Genomes Project reference data (Phase 3, release 5) ^29^. We then excluded entries whose RAF deviated more than 15% from the reference. This chosen threshold is subjective and was based on clear differentiation between correct and likely flipped alleles on the two diagonals (see **Figure S2**) as noted frequently in GWAS meta-analyses quality control procedures ^30^. For each analyzed cancer type, we extracted overlapping GWAS hits in our genotype data and estimated pairwise LD (r^2^) using the available allele dosages of the corresponding controls. For pairwise correlated SNPs (r^2^>0.5) or SNPs with multiple entries, we kept the SNP with the younger publication date (and smaller P value, if necessary) and excluded the other (**Figure S2 & Table S2**). Finally, we weighted the allele dosages of risk SNPs of the risk increasing alleles with their reported log odds ratios and calculated PRS as their sum. Namely, for subject *j* in MGI the PRS was of the form PRS_*j*_ = ∑_*i*_*β*_*i*_*G*_*ij*_ where the sum extends over all included loci, *β*_*i*_ are the log odds ratios retrieved from the GWAS catalog for locus *i* and *G*_*ij*_ was the measured dosage data for the risk allele on locus *i* in subject *j*. This variable was created for each MGI participant and for each cancer separately.

### Statistical Analysis

For the current study, we initially explored 30 cancer traits that had matching entries in the GWAS catalog (**Table S3**), and restricted our analysis to 12 cancer traits with at least 5 risk SNPs detected in the GWAS catalog after filtering that had relatively larger samples sizes in MGI (namely N≥250 cases) (Table 2). Logistic regression was used for all genetic association analysis. Firth's bias reduction method was applied to all single SNP and PRS models to resolve the problem of separation in logistic regression (Logistf in R package “EHR”) ^31-33^, a common problem for binary or categorical outcome models when for a certain part of the covariate space there is only one observed value of the outcome which often leads to very large parameter estimates and standard errors. Firth’s bias-reduction ^34^ is a penalized likelihood method that reduces the bias in such situations by adding a penalty term to the likelihood.

**Table 2.**
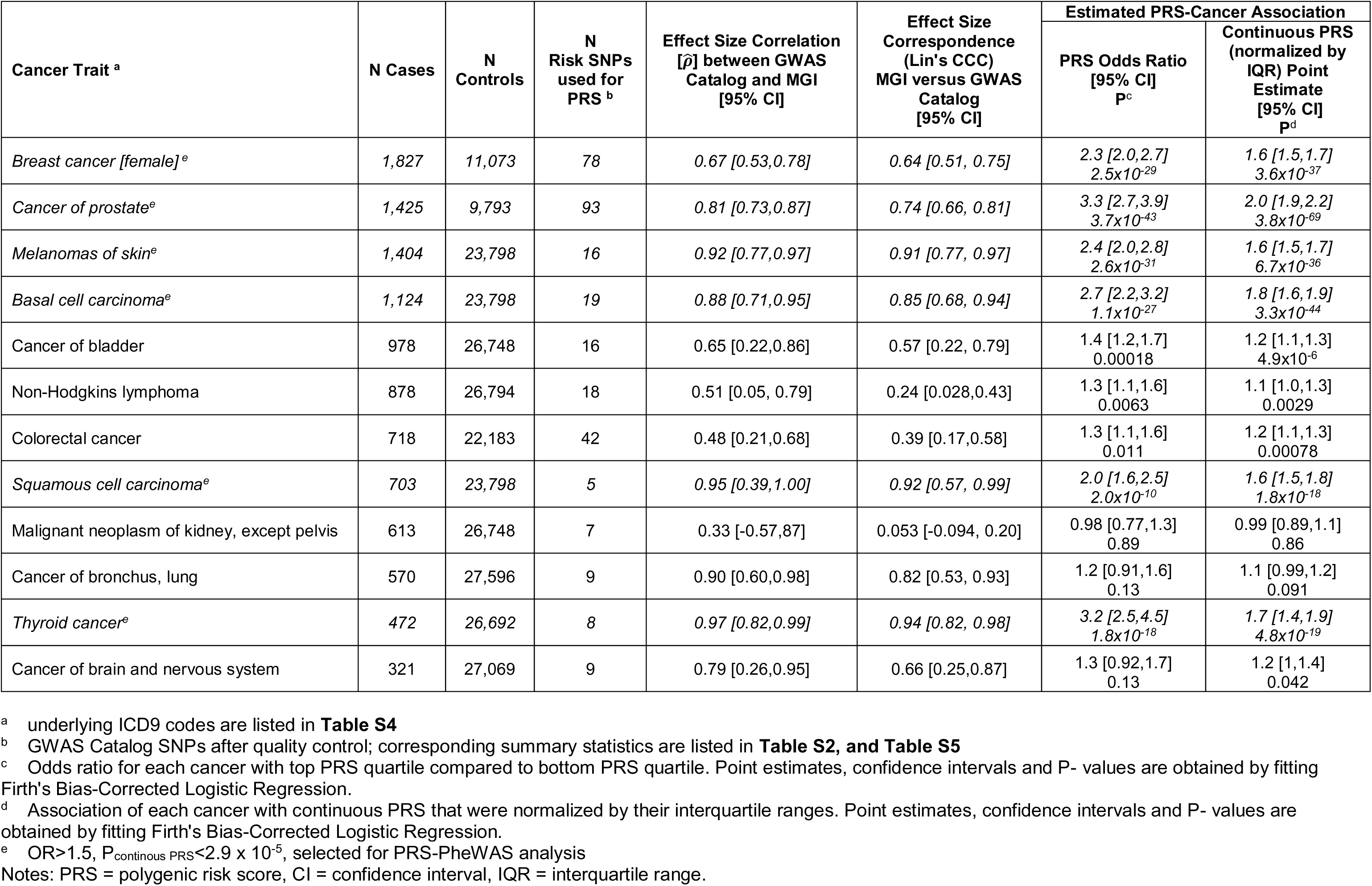
**Association analysis of cancer traits with at least five NHGRI EBI GWAS Catalog risk SNPs and more than 250 cases in MGI.**

To estimate the association of PRS with the primary cancer phenotype, we first determined the PRS quartiles using all control samples, categorized all samples according to these PRS quartiles and fitted Firth bias corrected logistic regression adjusting for age, sex, genotyping array, and the first four principal components. We report odds ratios corresponding to the top versus the bottom quartile PRS (reference), referred to as **PRS OR**. We also used continuous PRS instead of the categorized version as the covariate for enhanced power.

To compare reported associations of individual GWAS catalog SNPs with association observed in the MGI data set, we tested the association between reported GWAS hits and its corresponding trait using Firth bias-corrected logistic regression implemented in EPACTS (version 3.3, see **Web Resources**). Age, sex, genotyping array, and principal components 1–4 were included as covariates (see **Kinship and Ancestry Inference**). To determine the agreement of estimated effect sizes [estimated log(odds ratios)] between the MGI case-control studies and the published GWAS catalog hits, we estimated Pearson’s correlation coefficient [*ρ̂*] and Lin’s concordance measure between the two sets of coefficients^35; 36^. Towards more standard discovery type genome-wide association analysis with MGI data, we performed GWAS for the 9 cancer traits where the correspondence between the effect sizes were relatively strong [*ρ̂* ≥ 0.6], **Table 2**). For computationally efficient GWA analysis we used the score test-based saddle point approximation (SPA) ^37^ method adjusting for age, sex, genotyping array, the first four principal components. SPA was reported to provide accurate test statistics even for extremely unbalanced case-control ratios similar to Firth bias corrected logistic regression (see below) but was estimated to be 100 times faster than the latter ^37^.

For our primary PRS-PheWAS, for the six PRS in **Table 2** that showed strong and significant association, we conducted Firth bias-corrected logistic regression by fitting a model of the following form and repeated them for each of the 1,711 phenotypes. logit (P(Disease=1|PRS, Age, Sex, Array, PC))

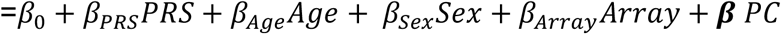

where the PCs were the first four principal components obtained from the principal component analysis of the genotyped GWAS markers and where “Array” represents the two genotyping array versions used in MGI accounting for potential batch effects. To adjust for multiple testing, we applied the conservative phenome-wide Bonferroni correction according to the 1,711 analyzed PheWAS codes (**Table S1**). Through a PheWAS plot, we present ‐log10 (*P*-values) corresponding to each of the 1,711 association tests for *H*_0_: *β*_*PRS*_ = 0. Directional arrows on the PheWAS plot indicate whether a phenome-wide significant trait was positively or negatively associated with the PRS.

Furthermore, our extensive sensitivity analyses included: (a) similar models adjusting for 20 PCs, (b) matching cases to control and conducting conditional logistic regression analysis and (c) using the unweighted risk allele counts as predictor. The reason for these three sensitivity analyses was to check (a) if the first 4 PCs were sufficient to control for population stratification, (b) if differences in age and sex distributions or extreme case-control ratios influenced the main analysis and (c) if ignoring effect sizes and using total risk allele count produced similar results. For (b), we matched cases and controls using the R package “MatchIt” and applied nearest neighbor matching for age, PC1-4 (using Mahalanobis-metric matching; matching window caliper/width of 0.25 standard deviations) and exact matching for sex. We considered a varying set of case:control matching ratios from 1:1 to 1:10. We observed gain in precision with increasing the number of matched controls per case, but the gain in precision became negligible after 1:10 matching ratio (**Table S6**). Moreover, for some cancers with large number of cases we could not attain 1:10 matching ratio for all cases and ended up with varying number of controls per case. For example, for prostate cancer the average number of matched controls per case was around 5 ^38^.

To investigate the possibility of the secondary trait associations with PRS being completely driven by the primary trait association, we performed a second set of PheWAS after excluding individuals affected with the primary cancer trait for which the PRS was constructed, referred to as “exclusion PRS PheWAS”. We applied “exclusion PRS PheWAS” instead of a PRS PheWAS that uses the primary cancer trait as covariate, because the control exclusion criteria implemented in the PheWAS phenotype construction pipeline will often eliminate these primary cancer cases from being eligible controls for some selected secondary related phenotypes and thus a logistic regression analysis will lead to complete separation^12^. We also stratified the MGI data set (or the corresponding gender subset depending on cancer type) into ten groups of equal size by PRS deciles and determined the percentage of observed cases for secondary traits in each risk decile and conducted a test of significance in difference in proportions across the deciles before and after removing individuals affected with cancer traits related to the primary cancer trait. As a follow-up tool to understand the secondary associations we created a plot to display the temporal ordering of diseases plotted against time of diagnoses. If not stated otherwise, analyses were performed using R 3.4.1 ^39^

## Results

In the current study, we report results obtained from 28,260 genotyped and unrelated samples of inferred European ancestry with available integrated ICD9-based EHR data. The study sample contains 53.5% females and the mean age is 54 years (see **Table 1 for summary**). We conducted our initial analysis on 12 cancer traits that after quality control had at least five independent risk variants in the NHGRI EBI GWAS Catalog and more than 250 cases in our cohort (**Table 2**, **Table S2**). **Table 2** summarizes data on 8,423 distinct individuals that were affected by at least one of the 12 cancers. Of these patients, 6,398 had one cancer, 1,574 had two cancers, and 451 had more than two cancer sites involved.

### Correspondence of MGI effect estimates with those reported in GWAS

To assess the calibration properties of the 12 ICD-9 based cancer case-control studies, we first compared the concordance of observed effect estimates (log odds ratios) from MGI with published effect estimates reported in the NHGRI EBI GWAS Catalog.

We found strong positive correlation (estimated Pearson’s correlation coefficient [*ρ̂*] > 0.6) between the MGI and GWAS reported estimates for 9 of the 12 cancers: female breast cancer (78 SNPs; [*ρ̂*]=0.67 [95% CI: 0.53,0.78]), prostate cancer (PCa; 93 SNPs; [*ρ̂*]=0.81 [0.73,0.87]), melanoma (16 SNPs; [*ρ̂*]=0.92 [0.77,0.97]), basal cell carcinoma (19 SNPs; [*ρ̂*]=0.88 [0.71,0.95]), bladder cancer (MIM: 109800; 16 SNPs; [*ρ̂*]=0.65 [0.22,0.86]), squamous cell carcinoma (5 SNPs; [*ρ̂*]=0.95 [0.39,1]), lung cancer (MIM: 211980; 9 SNPs; [*ρ̂*]=0.90 [0.6,0.98]), thyroid cancer (9 SNPs; [*ρ̂*]=0.79 [0.26,0.95]), and cancer of brain and nervous system (9 SNPs; [*ρ̂*]=0.79 [0.26,0.95]) (**Table 2**; **Table S5**; **Figure 1**; **Figure S3**).

**Figure 1.**
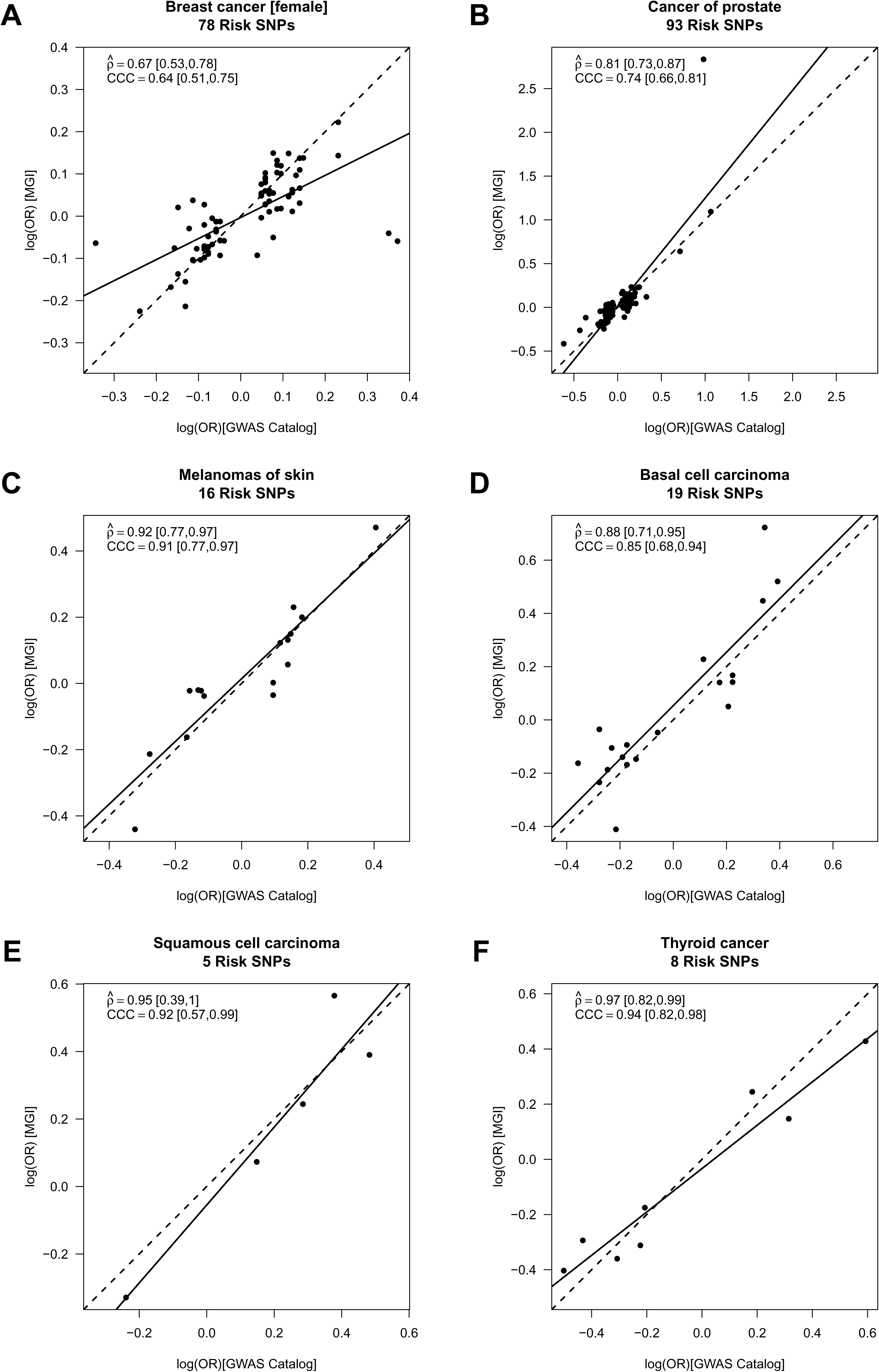
Calibration of association parameters. Calibration of association parameters between the MGI-GWAS and NHGRI-EBI GWAS Catalog derived effect estimates [log(OR)] for (**A**) breast cancer (females only), (**B**) cancer of prostate, (**C**) melanoma, (**D**) basal cell carcinoma, (**E**) squamous cell carcinoma, and (**F**) thyroid cancer. The agreement of two sets of SNP-specific beta coefficients (non-reference allele is the effect allele), their Pearson Correlation (Coefficient *ρ̂*, incl. 95% confidence interval and *P*) and Lin’s correspondence correlation (coefficient CCC; incl. 95% confidence interval) are shown; dashed line: perfect concordance; solid line: fitted line.

### Cancer GWAS in MGI

After having established strong positive correlation for 9 of the 12 cancer traits and thus a phenotype quality that appears to be in line with their corresponding published GWAS, we performed for each of these 9 cancers a GWAS to explore our ability to replicate and/or uncover cancer risk variants in a genome-wide setting. For the 9 cancers, we could replicate a total 55 of the 253 included risk SNPs with consistent effect orientation with P < 0.05 after correcting for the number of SNPs per phenotype (**Table S7**). We found genome-wide significant signals (P<5×10^-8^) for female breast cancer, melanoma of skin, basal cell carcinoma, squamous cell carcinoma and thyroid cancer. All but one of the genome-wide signals were found in loci already reported in the GWAS catalog for the corresponding cancer trait or related phenotypes. For instance, the four melanoma of skin loci with risk variants near *SLC45A2* (MIM 606202), *IRF4* (MIM 601900), *MC1R* (MIM 155555) and *ASIP/RALY* (MIM: 600201) were previously reported to be associated with melanoma, non-melanoma skin cancer, squamous cell carcinoma, or basal cell carcinoma ^40-44^. Also, the two breast cancer risk loci near *FGFR2* (MIM: 176943) and *FGF3/FGF4* (MIM: 164950 / 164980) as well as the thyroid cancer risk loci near *NRG1* (MIM: 142445) and *FOXE1* (MIM: 602617) were previously described ^45-48^ The only potentially novel finding was the SNP rs77909434 on chromosome 13 showing borderline genome-wide association with melanoma (MAF in cases = 5.3%; MAF in controls = 3.4%; P = 1.5×10^-8^) located 53 kb downstream of the Fibroblast Growth Factor 9 gene (*FGF9*, MIM: 600921) on chromosome 13. Since multiple phenotypes were involved in the genome-wide explorations, this SNP would not have passed the Bonferroni multiple testing correction for multiple GWAS. Further exploration in larger studies are warranted to substantiate this suggestive finding. We present GWAS Manhattan and QQ plots for all nine cancer traits in **Figure S4**.

Owing to the smaller sample sizes compared to the studies included in the NHGRI-EBI GWAS Catalog, only 8 of 253 catalog SNPs exceeded the genome-wide significance (**Table S5**). However, we found catalogued risk SNPs in **Table 2** were markedly enriched in the top 1% of GWAS associations, especially for the larger case/control studies. For example, 27 out of the 93 GWAS Catalog PCa risk SNPs fall in top 1% of associated SNPs in the MGI GWAS (with *P*<0.0083) (**Table S8**).

### Replicability of PRS Primary Cancer association

PRS integrates multiple SNPs, weighted by prior effect estimates and is expected to substantially improve the power to detect an association compared to an analysis with individual variants. To evaluate the association of PRS with the primary cancer trait, we estimated the OR for patients in the top risk quartile compared to the bottom quartile (PRS OR). Six of the 12 cancer PRS revealed an at least twofold enrichment of cases with P _Q1vsQ4_ < 2.0×10^-10;^ all of which also showed strong positive correlation between the MGI and GWAS reported estimates (see above): female breast cancer (PRS OR = 2.3 [95% CI: 2.0;2.7], P _Q1vsQ4_ = 2.5×10^-29^), prostate cancer (PRS OR = 3.3 [95% CI: 2.7;3.9], P _Q1vsQ4_ = 3.7×10^-43^), melanoma (PRS OR = 2.4 [95% CI: 2.0;2.8], P _Q1vsQ4_ = 2.6×10^-31^), basal cell carcinoma (PRS OR = 2.7 [95% CI: 2.2;3.2], P _Q1vsQ4_ = 1.1×10^-27^), squamous cell carcinoma (PRS OR = 2.0 [95% CI: 1.6;2.5], P _Q1vsQ4_ = 2.0×10^-10^), and thyroid cancer (PRS OR = 3.2 [95% CI: 2.5;4.5], P _Q1vsQ4_ = 1.8×10^-18^) (**Figure 1A-F**, **Table 2**).

The corresponding P-values obtained from Firth's bias-reduced logistic regression using continuous PRS were even stronger as expected and indicated that these six cancer traits would withstand a Bonferroni multiple testing correction in a phenome setting (1,711 traits; Pprs < 2.9 × 10^-5^): female breast cancer (P_PRS_=3.6×10^-37^), prostate cancer (P_PRS_=3.8×10^-69^), melanoma (P_PRS_=6.7×10^-36^), basal cell carcinoma (P_PRS_=3.3×10^-44^), cutaneous squamous cell carcinoma (SCC, P_PRS_=1.8×10^-18^), thyroid cancer (MIM: 188550) P_PRS_=4.8×10^-19^) (**Table 2**, **Table S3**). We excluded the remaining six cancer traits from further investigation, because in this initial analysis they showed only little or moderate association (PRS OR < 1.5) and consequently only modest power for subsequent exploration of phenome-wide associations (**Table 2**).

### PRS PheWAS

Next, we evaluated each of the six remaining PRS that were strongly associated with the primary cancer trait across a collection of 1,711 EHR-derived phenotypes (not limited to cancer traits) with at least 20 cases each (**Table S1**). For each of the six cancer PRS, we found strongest associations with their primary traits, except for squamous cell carcinoma PRS which revealed its strongest association with the more general skin cancer trait definition (P_PRS_ = 7.2×10^-61^) (**Figure 2**). Overall, we found no or little sign for inflation in our PheWAS results (median chi-squared based Lambda ≤ 1.16). Notably, we observed deflation for some PRS PheWAS that might be caused by lack of power especially for the phenotypes with small number of cases (**Figure S5**). We displayed the results from the three type of sensitivity analyses PheWAS next to the original results: conditional logistic regression results from 1:10 matched (for age, sex and first four PCs) case-control studies; adjusting for 20 principal components; using unweighted sum of risk allele counts instead of weighted PRS (**Figures S6-11**). The results remained robust with respect to these design and analytic choices.

**Figure 2.**
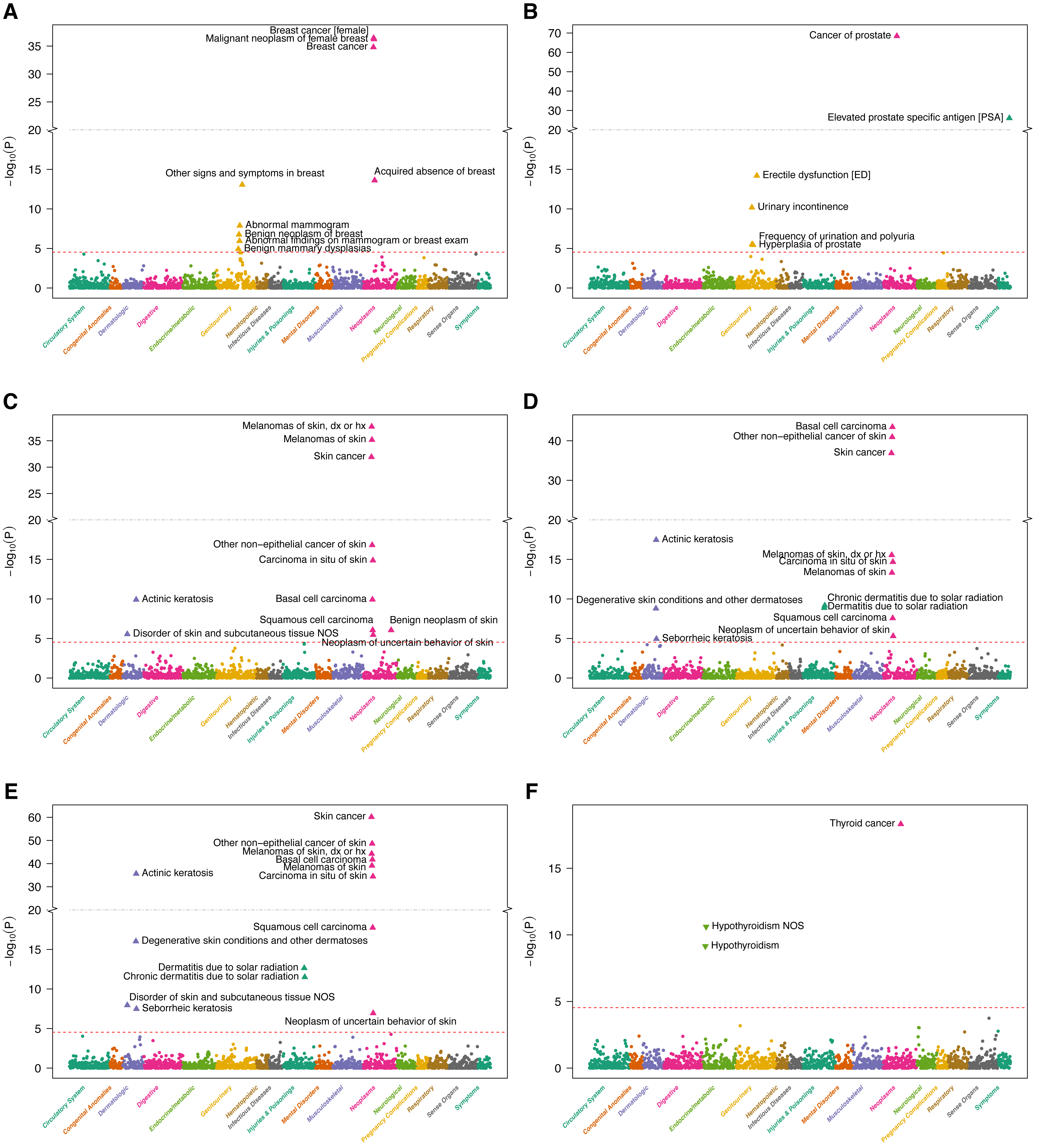
PRS PheWAS plots. PRS PheWAS plots for (**A**) breast cancer (females only), (**B**) cancer of prostate, (**C**) melanoma, (**D**) basal cell carcinoma, (**E**) squamous cell carcinoma, and (**F**) thyroid cancer. 1,711 traits are grouped into 16 color-coded categories as shown on the horizontal axis; the P-values for testing the associations of PRS with the traits are minus log-base-10-transformed and shown on the vertical axis. Triangles indicate phenome-wide significant associations with their effect orientation (up-pointing = risk increasing; down-pointing = risk decreasing). PRS upon multiplicity adjustment (see **Methods**). The solid horizontal line for *P*=2.9 ×10^-5^ cut-off.

### Secondary Associations

In addition, we identified for each PRS novel associations with secondary traits besides their primary traits (**Figure 2 A-F**, **Table S9**). For example, we observed associations of the three skin cancer PRSs (PRS for melanoma, basal cell carcinoma, and squamous cell carcinoma) with overall skin cancer and other skin cancer sub categories – expected due to their overlapping SNP sets – but also significantly associated with multiple dermatologic phenotypes, e.g. actinic keratosis (*P*_*PRS*_ < 1.2×10^-10^) and other degenerative skin conditions or disorders, all potential pre-cancer stages (**Figure 2C,D,E**).

Similarly, the female breast cancer PRS was associated not only with breast cancer (*P*_*PRS*_ =3.6×10^-37^) but also with acquired absence of breast (*P*_*PRS*_=2.4×10^-14^), abnormal mammogram (*P*_*PRS*_ =1.3×10^-8^), benign neoplasms of the breast (*P*_*PRS*_ =1.8×10^-7^) and benign mammary dysplasias (*P*_*PRS*_ =1.2×10^-5^) (**Figure 2A**). The PRS originally constructed for prostate cancer was associated with prostate cancer (*P*_*PRS*_ =3.8×10^-69^), as expected, but also with four additional traits: elevated prostate specific antigen (*P*_*PRS*_ =9.3×10^-27^), erectile dysfunction (*P*_*PRS*_ =6.3×10^-15^), urinary incontinence (*P*_*PRS*_ =6.6×10^-11^), frequency of urination and polyuria (*P*_*PRS*_ =2.9×10^-6^), and hyperplasia of prostate (*P*_*PRS*_ =3.6×10^-6^) (**Figure 2B**, **Table S9**).

While all of the above mentioned secondary trait associations were in the same effect orientation as their primary traits, i.e. increasing PRSs were associated with increased risk for the secondary trait, we observed an association of increasing thyroid cancer PRS with decreased risk for hypothyroidism (*P*_*PRS*_ =7.0×10^-10^) (**Figure 2F**).

### Exploring Secondary PRS PheWAS Associations via Exclusion PRS PheWAS

Since we already applied exclusion criteria to the controls during our phenome generation, e.g., individuals with elevated prostate specific antigen levels were excluded from being controls for prostate cancer and vice versa, we could not adjust for the primary cancer traits as a predictor in logistic regression models to identify independent secondary PRS PheWAS associations due to the issue of complete separation. To alternatively explore the secondary associations in PRS PheWAS (**Figure 2**), we proposed and performed “exclusion PRS PheWAS” by removing subjects affected with the cancer or related cancer traits for which the PRS was constructed. After removing all breast cancer cases (N = 1,894) no association with breast cancer PRS remained significant, e.g., acquired absence of breast (*P*_*PRS*_ = 0.52), abnormal mammogram (*P*_*PRS*_ = 0.76) or benign neoplasms of the breast (*P*_*PRS*_ = 0.49), indicating that the secondary trait associations were driven by the primary trait (**Figure S12A**). However, we noted that the majority of cases of the non-neoplasm phenotype “Acquired absence of breast” (>94.4%; 624 of 661) were removed in this step as they are highly correlated with breast cancer. We made similar observations for prostate cancer PRS where none of the previously detected secondary trait associations remained phenome-wide significant after removing all 1,425 prostate cancer cases (**Figure S12B**).

In contrast, we found a markedly stronger association between hypothyroidism and thyroid cancer PRS after removing 472 thyroid cancer cases (*P*_PRS_ = 4.7×10^-19^) compared to the full analysis (*P*_PRS_ = 7.0×10^-10^) which is consistent with the effect orientations between thyroid cancer PRS and hypothyroidism (**Figure 2F**).

To account for the substantial overlap between skin cancer sub types, e.g. 253 of the 1,404 individuals affected with melanoma are also affected by basal and/or squamous cell carcinoma (**Figure S13**) and to account for the likely intensified skin cancer screening of individuals that were diagnosed with skin cancer once in their life time, we excluded any type of skin cancer (N = 3,910) and repeated the PheWAS for melanoma, basal cell carcinoma, and squamous cell carcinoma PRS. After doing so, only actinic keratosis remained statistically associated with squamous cell carcinoma PRS while all of the previously observed associations mainly driven by skin cancer diagnoses disappeared (**Figure 2C-E** and **Figure S12C-E**). The association between squamous cell carcinoma PRS and actinic keratosis was less pronounced after excluding skin cancer cases but still remained phenome-wide significant (P_PRS_ = 2.3×10^-36^ versus *P*_PRS_ = 1.1 ×10^-12^).

To further understand the discovered secondary associations in the PRS PheWAS analyses (**Figure 2** and **Figure S12**), we conducted a simple follow-up analysis by stratifying the data into PRS deciles. We only discuss selected secondary trait associations for the prostate cancer (PCa), squamous cell carcinoma (SCC) and thyroid example in the main text and relegate their comprehensive analysis and a similar analysis of breast cancer, melanoma and basal cell carcinoma PRS to the supplemental material (**Table S10**). For prostate cancer, we stratified a total of 12,026 male individuals in MGI with age ≥30 years into deciles of PCa PRS. The observed PCa PRS associations in the PheWAS analysis are further supported by their respective increasing trait prevalences that are observed across 10 PCa PRS decile-stratified strata (**Figure 3A**; **Table S10**). These strata were not adjusted for confounders, but it is less likely that PRS is strongly associated with other covariates. A striking observation is that the proportion of PCa cases in lowest versus highest decile of PCa PRS is 5.4% versus 23.4% (Δ=18.0% [95% CI, 15.2 to 20.7%]; *P*=8.3×10^-36^) emphasizing that the PRS can distinguish well between high and low risk individuals in a realistic academic medical center population.

**Figure 3.**
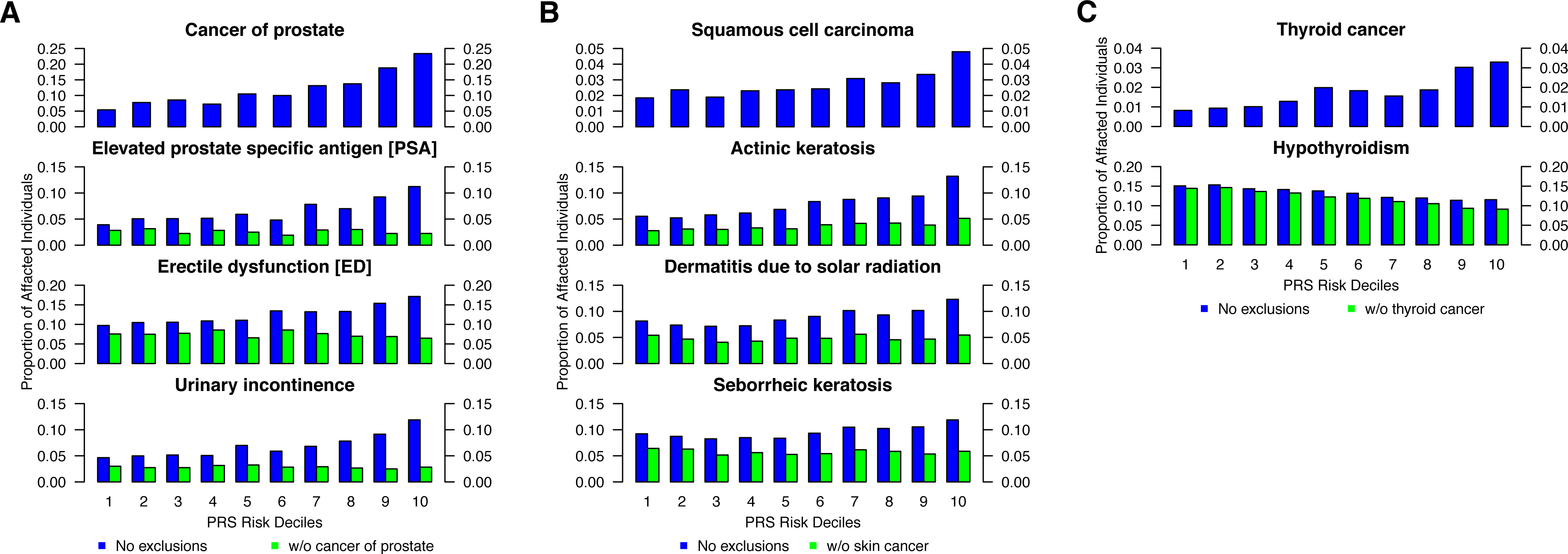
Proportion of primary and secondary traits stratified by PRS deciles. Percentage of primary and selected secondary traits in each cancer PRS category for (**A**) prostate cancer, (**B**) squamous cell carcinomas, and (**C**) thyroid cancer. Observed percentages in the MGI database as represented by the height of bars for each of 10 increasing decile-stratified PRS strata from left to right. The PRS’s underlying trait is shown on top and secondary traits below with (blue) and without (green) overlapping relevant cancer cases. Only individuals with age≥30 years were included in each analysis, with the prostate cancer PRS example only includes male individuals (see **Table S10** for detailed sample sizes and percentages).

Focusing on the secondary traits that reached phenome-wide significance with the PCa PRS, all these traits are known to be associated with PCa: erectile dysfunction (ED), and urinary incontinence (UI) – which commonly follows invasive surgical removal of the prostate – and elevated prostate specific antigen levels (ePSA) – which is a known biomarker for an increased PCa risk being closely monitored after prostatectomy. For example, when comparing the lowest versus the highest PRS risk decile, we found significant differences for ePSA (3.9% versus 11.2%; Δ=7.3% [95% CI, 5.1 to 9.5%]; *P*=2.0×10^-36^), ED (9.7% versus 17.1%; Δ=7.4% [95% CI, 4.6 to 10.2%]; *P*=1.4×10^-7^), and UI (4.7% versus 11.9%; Δ=7.2% [95% CI, 5.0 to 9.5%]; *P*=2.0×10^-10^). To test whether these associations are early indicator for PCa or whether they are driven by the fact that subjects affected by these secondary traits are also PCa cases (perhaps as a side effect of PCa treatment), we removed PCa cases and evaluated secondary disease prevalence across PCa PRS deciles. By doing so, prevalence of all secondary traits became roughly constant across PRS strata (**Figure 3A**; **Table S10**) and can be illustrated by the comparison of the proportions of the lowest versus the highest PRS risk decile: ePSA (2.8% versus 2.2%, Δ=-0.6% [95% CI, -1.9 to 0.8%]; *P*=0.44), ED (7.6% versus 6.5%, Δ=-1.1% [95% CI, -3.2 to 1.0%]; *P*=0.34), and UI (3.0% versus 2.8%, Δ=0.2% [95% CI, -1.6 to 1.3%]; *P*=0.90). Based on these observations, we hypothesize that the association of PCa PRS on the secondary traits ePSA, ED, and UI were driven by the PCa diagnosis, through either prior symptoms of PCa or prescribed medication, chemotherapy or surgical procedures for prostate removal (**Table S10**).

For SCC PRS stratification, there was a gradual increase of individuals affected with SCC with increasing PRS risk deciles, a trend that was also noted for actinic keratosis, dermatitis due to solar radiation, and seborrheic keratosis (**Figure 3B**). However, when excluding cases with skin cancer, the upward trend for the latter two phenotypes disappeared. The previously observed difference between the top and bottom PRS risk decile of individuals affected with actinic keratosis (5.5% versus 13.2% (Δ=7.7% [95% CI, 6.1 to 9.3%]; P=4.0×10^-21^) was markedly reduced after excluding skin cancer cases but still remained significant (2.8% versus 5.1% (Δ=2.4% [95% CI, 1.3 to 3.5%]; *P*=1.6×10^-5^) (**Table S10**) suggesting the potential for common genetic risk profiles between SCC and actinic keratosis. Since actinic keratosis is a known precursor for squamous cell carcinoma^49^, our approach indicated that it is possible to identify phenotypic risk factors through phenome-wide association scans and careful follow-up investigation of primary and secondary diagnoses.

Finally, we found an attenuated association between increasing thyroid cancer PRS and reduced risk for hypothyroidism: within all 25,681 samples >= 30 years of age the difference between bottom and top decile was Δ=-3.5% ([95% CI, -5.4 to -1.6%]; 15.1% versus 11.5%; *P*=2.5×10^-4^) and after excluding 452 thyroid cancer cases it increased to Δ=-5.3% ([95% CI, -7.1 to -3.5%]; 14.4% versus 9.1%; *P*=4.5×10^-9^) (**Table S10**). Several studies previously reported genetic overlap of a subset of thyroid cancer risk variants and variants associated with serum levels of thyroid stimulating hormone (TSH) which matches the current observed association between thyroid cancer risk and risk for hypothyroidism ^47; 48^.

To further our understanding of the observed secondary associations, we take advantage of the temporally resolved electronic health records data and explore the temporal order in which the diagnoses appear. **Figure 4** shows that actinic keratosis diagnosis mostly precedes the diagnosis of squamous cell carcinoma and sometimes by even 10 years. Erectile dysfunction or hypothyroidism, known side-effects of treatment of prostate and thyroid cancer (respectively), are mostly identified within a short timeframe of primary cancer diagnosis. Whereas elevated PSA, used as a screening tool for prostate cancer with known shared genetic correlation is observed mostly prior to a prostate cancer diagnosis and also after treatment as a prognostic marker. Having access to the electronic health records enables us to explore these temporally ordered data patterns and understand the explanation behind these secondary associations.

**Figure 4.**
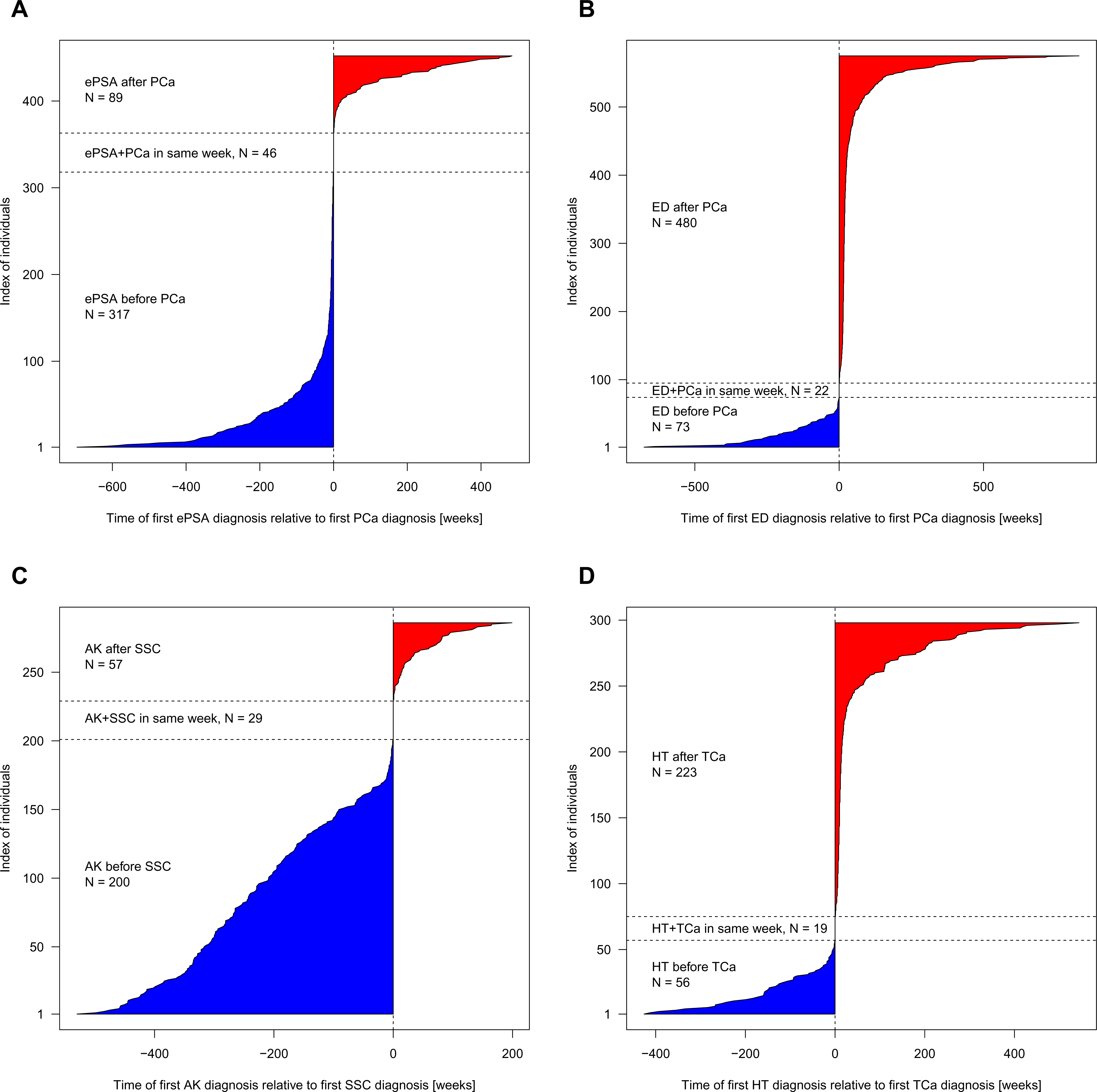
Temporal order of diagnoses: (**A**) elevated PSA levels (**ePSA**) and PCa in 452 individuals with PCa and ePSA; (**B**) erectile dysfunction (ED) and prostate cancer (PCa) in 575 individuals with ED and PCa; (**C**) actinic keratosis (AK) and squamous cell carcinoma (**SCC**) in 286 individuals with AK and SCC; and (**D**) hypothyrodism (HT) and thyroid cancer (TCa) in 298 individuals with HT and TC. The time of the first non-cancer diagnosis relative to the cancer diagnosis is shown in weeks; before (blue) and after (red) the cancer diagnosis.

## Discussion

Integration of large-scale biorepositories such as genetic data with EHRs are becoming increasingly common and indispensable for next-generation etiology studies. In this paper, we proposed, demonstrated and tested trait-specific PRS that summarize the results of large population-based GWAS studies towards cancer risk prediction in an actual academic medical center population managed by Michigan Medicine. Data repositories like MGI, allow us to explore many traits simultaneously whereas population-based case-control studies focus on one specific trait. It is indeed encouraging that the results of population-based studies corroborate with the phenotypes computed from EHR data. We found improved trait prediction power of the composite PRS compared to single-SNP analyses. We also replicated catalogued associations of SNPs for some cancer traits, observed excellent correspondence of effect estimates and discovered novel secondary trait associations with cancer PRS that were not driven by the primary cancer diagnoses. To our knowledge this is the first comprehensive PRS PheWAS study and the first PheWAS study focused on cancer.

We have introduced several novel analytic strategies in this paper. We presented a principled framework and quality-control pipeline to create a PRS from a large curated, public database and to perform PRS PheWAS in a potentially biased sample. We introduced a primary PheWAS using Firth's bias-reduced logistic regression which has the advantage of resolving the problem of separation in logistic regression and providing well-controlled type I error rates for unbalanced case control studies with relatively small sample counts ^31; 32; 50^. These issues are often present in large EHR-based phenomes where controls are frequently hundredfold more abundant than cases. In addition, we conducted thorough sensitivity analyses to check the robustness of our findings by using PheWAS with unweighted risk allele counts, adjusting for 20 PCs and PRS PheWAS based on matched controls. All our reported results remained robust under these sensitivity analyses.

To distinguish PRS-trait associations that truly derive from a shared genetic risk profile from secondary associations that are potentially driven by the primary trait (for example urinary incontinence or erectile dysfunction following prostate cancer treatment), we further introduced a modified PRS PheWAS approach that excludes the PRS’s underlying cancer traits. While reducing overall sample size, this “exclusion PRS PheWAS” approach is statistically preferable in contrast to a PRS PheWAS that conditions on the primary cancer trait. A conditional PheWAS approach is often affected by unilaterally applied exclusion criteria of controls that occur during the phenome construction, e.g. PCa cases were excluded from being eligible controls for elevated PSA levels and vice versa. Our approach could directly discard trait associations driven by the primary cancer diagnosis and has the potential to identify clinically useful diagnostic traits among many that are conveniently measured in panel tests of biomarkers. When an association with a secondary trait disappears by removing the primary cancer cases in an exclusion PheWAS, there can be several alternative explanations: truly shared genetic correlation, intensified screening/examination due to detection of an initial cancer, a screening biomarker/pre-cancer phenotype or simply post treatment effects. We used the temporal ordering of the diagnoses to understand which of the above explanations appear plausible for a given scenario. Further exploration of our findings in larger biobank studies, like the UK Biobank study, is warranted and will empower a deeper understanding of relevant pre-cancer traits ^51^.

There are several limitations to the current study. We decided to rely on the associations reported in the NHGRI-EBI GWAS Catalog instead of focusing on the latest and largest GWAS study specific for each cancer trait. Our rationale for choosing the NHGRI-EBI GWAS Catalog as our source for extracting summary statistics were primarily three-fold: (1) <u>Data quality</u>: Summary statistics in the GWAS Catalog underwent a detailed expert curation and harmonization ^28; 52^ that avoids redundancy, allows reliable SNP position extraction, and most importantly ancestry matching; We wanted to use a database that is publicly accessible and applies the same set of criteria to update reported results across a wide variety of phenotypes. (2) <u>Reproducibility</u>: We provided detailed instructions on how to extract and filter GWAS Catalog summary statistics to construct PRS. This will allow interested readers to easily apply our approach to the regularly updated GWAS catalog versions or to a different ancestry group and/or broad set of disease categories without requiring detailed and deep literature searches that could be somewhat subjective. (3) <u>Scalability to Multiple Phenotypes:</u> One can construct PRS for specific cancers of primary interest from the latest GWAS meta-analyses following the same prescriptions we provided. Using the latest GWAS result is likely to enhance power of a PRS PheWAS. Similarly using a PRS that is based on a truly polygenic model with many more variant (or the entire genome) instead of considering the GWAS hits may reveal new associations.

We restricted our analysis to GWAS results from studies of broad European ancestry to match them to our cohort of predominantly European ancestry and to allow an extra filtering of potential swaps in directionality of risk allele in published GWAS studies that otherwise could have negatively affected the correlative properties of our constructed PRS. One could modify or extend construction of PRS based on global ancestry, functionality of the variant and use other weighting schemes. Stratifying the present analysis by young onset cancers, metastatic/aggressive cancers or tumor subtype will shed further insight into cancer biology, cancer genetics and specificity of the PRS-cancer association. We have mostly ignored the temporal ordering in the diagnoses codes by defining dichotomous phenotypes of interest. Exploring the time-stamped data in greater detail may be instrumental in understanding the secondary associations like the negative association between hypothyroidism and thyroid cancer PRS.

Though we note some very encouraging and promising results for the cancer traits with modest number of cases and controls and with a larger number of variants reported in the NHGRI-EBI GWAS catalog, we also note that the correlation of effect estimates or the PRS-cancer association was not very strong for some cancers (**Table 2**). This could be due to limited sample size/power, heterogeneity in the definition of the cancer phenotype, incomplete specification of PRS, differences in allele frequencies in the MGI population, or misclassification of ICD-9 codes. To address concerns with misclassification we conducted detailed chart review of 50 randomly sampled cases with at least one cancer PheWAS code and verified their primary and secondary cancer diagnosis. We could verify 149 of the 151 diagnoses and found 49/50 patients to have accurate record of their cancer diagnosis. Based on this we conclude that the rate of misclassification will likely be low for ICD 9 codes associated with cancer.

In this paper, we have focused on cancer traits. The low misclassification rate of cancer traits, typical within academic health and cancer centers, along with effective sensitivity analyses partly protect the results against imprecise case definitions and confounding. For non-cancer disease traits, more stringent ICD-9 defined cases, e.g., by repeated ICD-9 diagnoses, of adequate sample sizes might alleviate the biases from case misclassification. Future analysis will need to control for potentially different levels of misclassification error across phenotypes.

Our phenome comprised a total of 1,711 ICD9-based phenotypes and by its implemented design of hierarchical phenotypes with different levels of specificity induce a certain degree of redundancy. While we applied the multiple testing correction for 1,711 performed tests, we acknowledge that this threshold might be too conservative. For examples, we estimated a maximal set of 1,452 phenotypes with all pair-wise correlations r^2^ < 0.5 before applying any exclusion criteria to the controls. In addition, the PheWAS approach often applies similar exclusion criteria to related phenotypes and thereby further reduces the observable independence of case-control studies. Future studies are needed to determine the effective number of independent tests in such a phenome-wide analysis. Besides the ICD-9 codes used for case and control definitions, EHR databases generally contain vast amount of additional patient information including ICD-10 codes, temporal laboratory tests, drug prescriptions, inpatient and outpatient records, etc. Future analyses that leverage these heterogeneous data sources that might be predictive of disease outcomes could further improve disease risk predictions. It will be interesting to study whether PRS for cancer risk behaviors like smoking, alcohol and obesity predict cancer phenotypes. Tailored and validated models capable of integrating multiple sources of molecular and environmental data data for predicting risks of disease will be crucial.

## Supplemental Data

Supplemental Data include 13 figures and 10 tables.

## Acknowledgments

The authors acknowledge the University of Michigan Medical School Central Biorepository for providing biospecimen storage, management, and distribution services in support of the research reported in this publication.

## Web Resources

Michigan Genomics Initiative, https://www.michigangenomics.org

OMIM, http://www.omim.org

BAF Regress, http://genome.sph.umich.edu/wiki/BAFRegress

PLINK v1.90, https://www.cog-genomics.org/plink2

KING v2.0, http://people.virginia.edu/~wc9c/KING

Human Genome Diversity Project reference panel, http://csg.sph.umich.edu/chaolong/LASER

PheWAS R package, https://github.com/PheWAS/PheWAS

NHGRI-EBI GWAS Catalog, https://www.ebi.ac.uk/gwas

UCSC genome browser, http://genome.ucsc.edu

EPACTS, http://genome.sph.umich.edu/wiki

